# *Salmon* provides accurate, fast, and bias-aware transcript expression estimates using dual-phase inference

**DOI:** 10.1101/021592

**Authors:** Rob Patro, Geet Duggal, Michael I Love, Rafael A Irizarry, Carl Kingsford

**Author notes:** work done while GD was at CMU.

## Abstract

We introduce *Salmon*, a new method for quantifying transcript abundance from RNA-seq reads that is highly-accurate and very fast. *Salmon* is the first transcriptome-wide quantifier to model and correct for fragment GC content bias, which we demonstrate substantially improves the accuracy of abundance estimates and the reliability of subsequent differential expression analysis compared to existing methods that do not account for these biases. *Salmon* achieves its speed and accuracy by combining a new dual-phase parallel inference algorithm and feature-rich bias models with an ultra-fast read mapping procedure. These innovations yield both exceptional accuracy and order-of-magnitude speed benefits over alignment-based methods.

---

Estimating transcript abundance across cell types, species, and conditions is a fundamental task in genomics. For example, these estimates are used for the classification of diseases and their subtypes [1], for understanding expression changes during development [2], and tracking the progression of cancer [3]. Accurate and efficient quantification of transcript abundance from RNA-seq data is an especially pressing problem due to both the wide range of technical biases that affect the RNA-seq fragmentation, amplification and sequencing process [4, 5] and the exponentially increasing number of experiments and the growing adoption of expression data for medical diagnosis [6]. Traditional quantification algorithms — those that make use of full alignments of the sequencing reads to the genome or transcriptome to compute abundances — require significant computational resources [7] and do not scale well with the rate at which data is produced [8]. Addressing the efficiency problem has been the focus of much recent work in the area of transcript-level quantification. For example, the quantification tool *Sailfish* [9] achieved an order of magnitude speed improvement over previous approaches by replacing traditional read alignment with the allocation of exact k-mers to transcripts but can sometimes produce less accurate estimates for paired-end data or for stranded protocols. The recently-introduced quantification tool, *kallisto* [10], achieves similar speed improvements and further reduces the gap in accuracy with traditional alignment-based methods by replacing the k-mer counting approach used in *Sailfish* with a procedure called pseudoalignment. Unlike pseudoalignment, *Salmon*’s lightweight mapping procedure tracks the position and orientation of all mapped fragments, and in conjunction with the abundances learned in the online phase of the inference algorithm, makes use of per-fragment conditional probabilities to estimate auxiliary models, bias terms, and aggregate weights for its rich equivalence classes.

However, existing methods for transcriptome-wide abundance estimation, including both traditional, alignment-based approaches and the recently-introduced ultra-fast methods, lack sample-specific bias models rich enough to capture many of the important effects, like fragment GC content bias, that are observed in experimental data and that can lead to, for example, unacceptable false positive rates in differential expression studies [5].

Our novel quantification procedure called *Salmon* (Supplementary Fig. 1) achieves best-in-class accuracy, employs high-fidelity, sample-specific bias models, and simultaneously achieves the same order-of-magnitude speed benefits as *kallisto* and *Sailfish*. Using experimental data from the GEUVADIS [11] and SEQC [28] studies as well as synthetic data from both the *Polyester* [13] and *RSEM-sim* [14] simulators, we benchmark *Salmon* against both *kallisto* [10] (as representative of a state-of-the-art alignment-free method) and *eXpress* [15] + *Bowtie2* [16] (as representative of a state-of-the art alignment-based method); both of these methods also implement their own bias models. We show that *Salmon* typically outperforms both *kallisto* and *eXpress* in terms of accuracy (Fig. 1a-d, Supplementary Fig. 4), often by a substantial margin.

**Figure 1:**
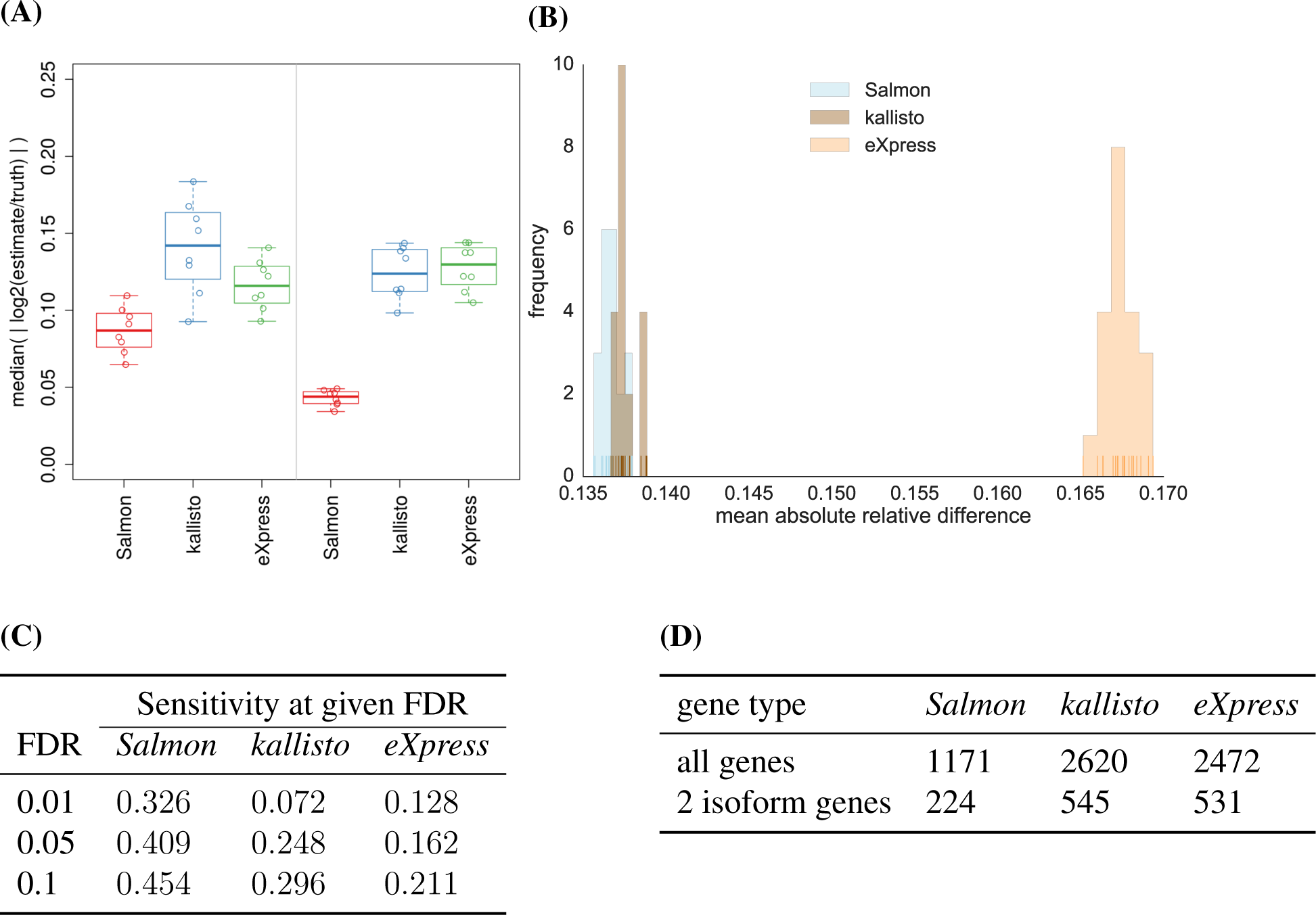
(A) The median of absolute log fold changes (lfc) between the estimated and true abundances under all 16 replicates of the *Polyester* simulated data. The closer the lfc to zero, the more similar the true and estimated abundances. The left and right panels show the distribution of the log fold changes under samples simulated with different GC-bias curves learned from experimental data (details in **Online methods, Ground truth simulated data**). (B) The distribution of mean absolute relative differences (MARDs), as described in **Online methods, Metrics for accuracy**, of *Salmon*, *kallisto* and *eXpress* under 20 simulated replicates generated by *RSEM-sim*. *Salmon* and *kallisto* yeild similar MARDs, though *Salmon*’s distribution of MARDs is significantly smaller (Mann-Whitney *U* test, *p* = 0.00017) than those of *kallisto*. Both methods outperform *eXpress* (Mann-Whitney *U* test, *p* = 3.39781 × 10^−8^). (C) At typical FDR values, the sensitivity of finding truly DE transcripts using *Salmon*’s estimates is 53% − 450% greater than that using *kallisto*’s estimates and 210% − 250% greater than that using *eXpress*’ estimates for the *Polyester* simulated data. (D) For 30 GEUVADIS samples, the number of transcripts called as DE at an expected FDR of 1% when the contrast between groups is simply a technical confound (i.e. the center at which they were sequenced). *Salmon* produces fewer than half as many DE calls as the other methods.

Further, *Salmon*’s dual-phase inference algorithm and rich bias models yield considerably improved inter-replicate concordance (Supplementary Fig. 2) compared to both *kallisto* and *eXpress*. For example, when used for differential expression (DE) testing, the quantification estimates produced by *Salmon* exhibit markedly higher sensitivity at the same false discovery rate than those produced by *kallisto* or *eXpress* (Table 1C) — achieving a sensitivity 53% to 250% higher, at the same FDRs, compared with existing methods. Likewise, *Salmon* produces fewer false-positive differential expression calls in comparisons that are expected to contain few true differences in transcript expression (Table 1D). These benefits of *Salmon* over other methods persist in gene-level analysis as well, where the use of *Salmon*’s estimates for gene-level DE analysis leads to a decrease by a factor of ∼ 2.6 in the number of genes that are called as DE (Supplementary Table 1). Supplementary Fig. 5 shows specific examples from the GEUVADIS experiments where dominant isoform switching is observed (*p* < 1 × 10^−6^) between samples under the quantification estimates produced by *kallisto* or *eXpress*, but this isoform switching is eliminated under the abundance estimates of *Salmon* that account for fragment GC bias. In idealized simulations, like those generated by *RSEM-sim*, where realistic biases are not simulated, the accuracy estimates of different methods tend to be more similar to one another (Fig. 1b, Supplementary Fig. 6). These idealized *RSEM-sim* simulation results, where the fragments are generated without bias and in perfect accordance with the generative model adopted by the quantifiers, serve as a useful measure of the internal consistency of the algorithms and have been used in other validation contexts [14, 10]. However, we expect our results on the SEQC [28], GEUVADIS [11], and *Polyester* [13] (simulated with bias) data sets to be more representative of typical real-world performance.

*Salmon* incorporates a rich model of experimental biases, which allows it to account for the effects of sample-specific parameters and biases that are typical of RNA-seq data, including positional biases in coverage, sequence-specific biases at both the 5′ and 3′ end of sequenced fragments, fragment-level GC bias, strand-specific protocols, and the fragment length distribution. These biases are automatically learned in the online phase of the algorithm, and are encoded in a fragment-transcript agreement model (**Online methods, Fragment-transcript agreement model**). In this model, fragment-transcript assignment scores are defined as proportional to (1) the chance of observing a fragment length given a particular transcript/isoform of a gene, (2) the chance that a fragment starts at a particular position on the transcript, (3) the concordance of the fragment aligning with a user-defined sequencing library format (e.g. a paired ended, stranded protocol), and (4) the chance that the fragment came from the transcript based on a score obtained from the the alignment procedure (if alignments are being used). Additionally, based on the computed mappings, *Salmon* gathers information about the positional, sequence-specific and fragment GC content of the observed fragments. *Salmon* incorporates these biases and experimental parameters by learning auxiliary models that describe the relevant distributions and maintaining ‘rich equivalence classes’ of fragments (**Online methods, Equivalence classes**) that act as an efficient representation of the sequenced fragments during the offline inference phase and speed up the process of estimating transcript abundances.

*Salmon*’s two-phase, parallel inference procedure consists of both a streaming, online inference phase, where estimates of transcript abundance are continuously updated after considering each small batch of reads, and an offline inference phase that operates over a highly-reduced representation of the sequencing experiment (**Online methods, Online phase** and **Offline phase**; Illustration of method in Supplementary Fig. 1). This two-phase inference procedure allows *Salmon* to build a probabilsitic model of the sequencing experiment that incorporates information not considered by *Sailfish* [9] and *kallisto* [10]. During the online inference phase, *Salmon* learns and continuously updates transcript-level abundance estimates. In turn, this allows the evaluation of per-fragment probabilities that are not directly represented in the factorized likelihood function (**Offline phase**). These probabilities enable accounting, proportionally, for all of the potential mapping locations of a fragment when estimating experiment and bias model parameters.

*Salmon* is designed take advantage of multiple CPU cores, and the mapping and inference procedures scale well with the number of reads in an experiment. *Salmon* can quantify abundance either via a builtin, ultra-fast read-mapping procedure [17], or using pre-computed alignments provided in SAM or BAM format. This approach allows *Salmon* to acheive high accuracy while maintaining a speed similar to that of *kallisto* [10]. For example, *Salmon* can quantify a data set of approximately 600 million reads (75bp, paired-end) in 23 minutes using 30 threads — this roughly matches the speed of the recently-introduced *kallisto*, which took 20 minutes to complete the same task (wall clock time on a 24 core (with hyperthreading) machine with 256Gb of RAM. Each core is Intel R ^®^Xeon^®^ CPU (E5-4607 v2 2.60GHz).

The simultaneous speed and accuracy of *Salmon* is achieved using the dual-phase approach described above together with quasi-mapping [17]. Therefore, *Salmon* encompasses both the “alignment” and “quantification” phases that are required by more traditional quantification pipelines in a single tool. A quasi-mapping represents a match between a sequenced fragment and a transcript and consists of a tuple *m_i_* = (*t, p_ℓ_, p_r_, o_ℓ_, o_r_*) containing the transcript *t* to which the fragment maps, the positions *p_ℓ_, p_r_* where the left and right ends of the fragment map, and the orientations *o_ℓ_, o_r_* with which the fragment ends map, but not the nucleotide-to-nucleotide correspondence between the fragment and transcript. When run in quasi-mapping mode, *Salmon* takes as input an index of the transcriptome and a set of *raw* sequencing reads (i.e. unaligned reads in FASTA/Q format) and performs quantification directly without generating any intermediate alignment files. This saves considerable time and space, since quasi-mapping is considerably faster than traditional alignment. Thus, while *Salmon* is also capable of performing quantification using existing alignments of a sequencing experiment to the transcriptome, if a users prefers to provide these, we anticipate that *Salmon* will be primarily used in quasi-mapping mode.

*Salmon*’s approach is unique in the way that it combines useful models of experimental data with an efficient parallel inference procedure. This combination has produced some of the most accurate expression estimates to date without sacrificing the order of magnitude speed improvements enjoyed by recent approaches (e.g. [9], [10]). *Salmon*’s ability to compute high-quality estimates of transcript abundances at the scale of thousands of samples, while also accounting for the prevalent technical biases affecting transcript quantification [18], will enable individual expression experiments to be interpreted in the context of many rapidly growing sequence expression databases. This will allow for a more comprehensive comparison of the similarity of experiments across large populations of individuals and across different environmental conditions and cell types. *Salmon* is open-source and freely-licensed (GPLv3). It is written in C++11, and is available at https://github.com/COMBINE-lab/salmon.

## Acknowledgements

This research is funded in part by the Gordon and Betty Moore Foundation’s Data- Driven Discovery Initiative through Grant GBMF4554 to C.K. It is partially funded by the US National Science Foundation (CCF-1256087, CCF-1319998, BBSRC-NSF/BIO-1564917) and the US National Institutes of Health (R21HG006913, R01HG007104). C.K. received support as an Alfred P. Sloan Research Fellow. This work was partially completed while G.D. was a postdoctoral fellow in the Computational Biology Department at Carnegie Mellon University. M.L. was supported by NIH grant 5T32CA009337-35. R.I. was supported by NIH R01 grant HG005220.

### Author Contributions

R.P. and C.K. designed the method, which was implemented by R.P. R.P., G.D., M.L., R.I. and C.K. designed the experiments and R.P., G.D. and M.L. conducted the experiments. R.P., G.D., M.L., R.I. and C.K. wrote the manuscript.

## Online methods

### Objectives and models for abundance estimation

Our main goal is to quantify, given a known transcriptome 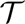 and a set of sequenced fragments *ℱ*, the relative abundance of each transcript in our input sample. This problem is challenging both statistically and computationally. The main statistical challenges derive from need to resolve a complex and often very high-dimensional mixture model (i.e. estimating the relative abundances of the transcripts given the collection of ambiguously mapping sequenced fragments). The main computational challenges derive from the need to process datasets that commonly consist of tens of millions of fragments, in conditions where each fragment might reasonably map to many different transcripts. We lay out below how we tackle these challenges, beginning with a description of our assumed generative model of the sequencing experiment, upon which we will perform inference to estimate transcript abundances.

We also make note of the notation we use in the methods described below. Here, we use the vertical bar *|* to indicate that the fixed quantities following are parameters used to calculate the probability. For the Bayesian objective, the notation implies conditioning on these random variables.

#### Generative process

Assume that, for a particular sequencing experiment, the underlying true transcriptome is given as 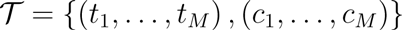, where each *t_i_* is the nucleotide sequence of some transcript (an isoform of some gene) and each *c_i_* is the corresponding number of copies of *t_i_* in the sample. Further, we denote by *ℓ_i_* the length of transcript *t_i_* and by 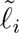 the *effective length* of transcript *t_i_*, as defined in Equation (1). We adopt a generative model of the sequencing experiment that dictates that, in the absence of experimental bias, library fragments are sampled proportional to 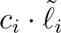. That is, the probability of drawing a sequencing fragment from some position on a particular transcript *t_i_* is proportional the total fraction of all nucleotides in the sample that originate from a copy of *t_i_*. This quantity is called the nucleotide fraction [14]:

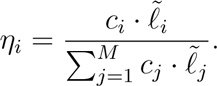

The true nucleotide fractions, *η*, though not directly observable, would provide us with a way to measure the true relative abundance of each transcript in our sample. Specifically, if we normalize the *η_i_* by the effective transcript length 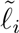, we obtain a quantity

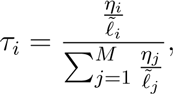

called the transcript fraction [14]. These *τ* can be used to immediately compute common measures of relative transcript abundance like transcripts per million (TPM). The TPM measure for a particular transcript is the number of copies of this transcript that we would expect to exist in a collection of one million transcripts, assuming this collection had exactly same distribution of abundances as our sample. The TPM for transcript *t_i_*, is given by TPM_*i*_ = *τ_i_*10^6^. Of course, in a real sequencing experiment, there are numerous biases and sampling effects that may alter the above assumptions, and accounting for them is essential for accurate inference. Below we describe how *Salmon* accounts for 5’ and 3’ sequence-specific biases (which are not considered separately by *kallisto*) and fragment GC bias which is modeled by neither *kallisto* nor *eXpress*.

#### Effective length

A transcript’s effective length depends on the empirical fragment length distribution of the underlying sample and the length of the transcript. It accounts for the fact that the range of fragment sizes that can be sampled is limited near the ends of a transcript. Here, fragments refer to the (potentially size-selected) cDNA fragments of the underlying library, from the ends of which sequencing reads are generated. In paired-end data, the mapping positions of the reads can be used to infer the empirical distribution of fragment lengths in the underlying library, while the expected mean and standard deviation of this distribution must be provided for single-end libraries. We compute the effective transcript lengths using the approach of [10], which defines the effective length of a transcript *t_i_* as

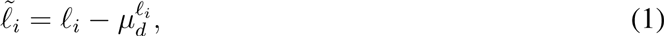

where 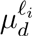 is the mean of the truncated empirical fragment length distribution. Specifically, let *d* be the empirical fragment length distribution, and Pr {*X* = *x*} be the probability of drawing a fragment of length *x* under *d*, then 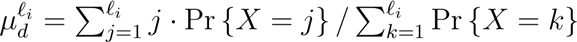.

Given a collection of observations (raw sequenced fragments or alignments thereof), and a model similar to the one described above, there are numerous approaches to inferring the relative abundance of the transcripts in the target transcriptome, 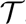. Here we describe two basic inference schemes, both available in *Salmon*, which are commonly used to perform inference in such a model. All of the results reported in the manuscript were computed using the maximum likelihood objective (i.e. the EM algorithm) in the offline phase, which is the default in *Salmon*.

### Maximum likelihood objective

The first scheme takes a maximum likelihood approach to solving for the quantities of interest. Specifically, if we assume that all fragments are generated independently, and we are given a vector of known nucleotide fractions *η*, a binary matrix of transcript-fragment assignment *Z* where *z_ji_* = 1 if fragment *j* is derived from transcript *i*, and the set of transcripts 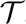, we can write the probability of observing a set of sequenced fragments *ℱ* as:

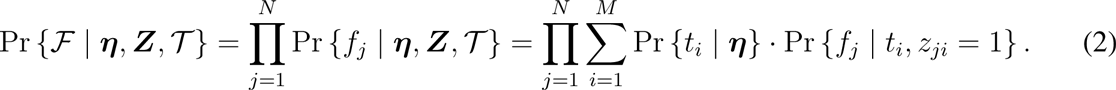

Where |*ℱ*| = *N* is the number of sequenced fragments, Pr {*t_i_*|*η*} is the probability of selecting transcript *t_i_* to generate some fragment given the nucleotide fraction *η*, and we have that Pr {*t_i_*|*η*} = *η_i_*. Pr {*f_j_*|*t_i_, z_ji_* = 1} is the probability of generating fragment *j* given that it came from transcript *i*. We will use Pr{*f_j_*|*t_i_*} as shorthand for Pr {*f_j_*|*t_i_, z_ji_* = 1} since Pr {*f_j_|t_i_, z_ji_* = 0} is uniformly 0. The determination of Pr {*f_j_|t_i_*} is defined in further detail in **Fragment-transcript agreement model**. The likelihood associated with this objective can be optimized using the EM algorithm as in [14].

### Bayesian objective

One can also take a Bayesian approach to transcript abundance inference as done in [19, 20]. In this approach, rather than directly seeking maximum likelihood estimates of the parameters of interest, we want to infer the posterior distribution of η. In the notation of [19], we wish to infer 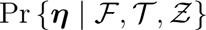 — the posterior distribution of nucleotide fractions given the transcriptome 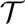 and the observed fragments *ℱ*. This distribution can be written as:

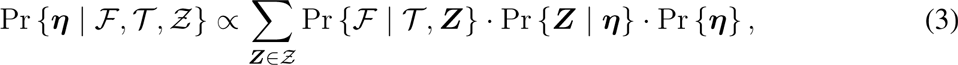

where

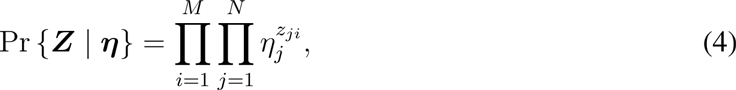

and

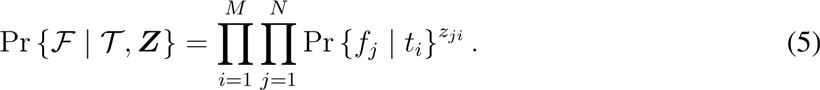

Unfortunately, direct inference on the distribution 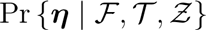 is intractable because its evaluation requires the summation over the exponentially large latent variable configuration space 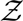. Since the posterior distribution cannot be directly estimated, we must rely on some form of approximate inference. One particularly attractive approach is to apply variational Bayesian (VB) inference in which some tractable approximation to the posterior distribution is assumed.

Subsequently, one seeks the parameters for the approximate posterior under which it best matches the true posterior. Essentially, this turns the inference problem into an optimization problem — finding the optimal set of parameters — which can be efficiently solved by a number of different algorithms. In particular, variational inference seeks to find the parameters for the approximate posterior that minimizes the Kullback-Leibler (KL) divergence between the approximate and true posterior distribution. Though the true posterior may be intractable, this minimization can be achieved by maximizing a lower-bound on the marginal likelihood of the posterior distribution [21], written in terms of the approximate posterior. *Salmon* optimizes the collapsed variational Bayesian objective [19] in its online phase and the full variational Bayesian objective [20] in the variational Bayesian mode of its offline phase (see **Offline phase**).

### Fragment-transcript agreement model

We model the conditional probability Pr {*f_j_|t_i_*} for generating *f_j_* given *t_i_* using a number of auxiliary terms. These terms come from auxiliary models whose parameters do not explicitly depend upon the current estimates of transcript abundances. Thus, once the parameters of these these models have been learned and are fixed, these terms do not change even when the estimate for Pr {*t_i_|η*} = *η_i_* needs to be updated. *Salmon* uses the following auxiliary terms:

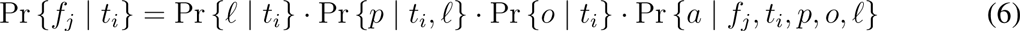

Where Pr {*ℓ|t_i_*} is the probability of drawing a fragment of the inferred length, *ℓ*, given *t_i_*, and is evaluated based on an observed empirical fragment length distribution. Pr {*p|t_i_, ℓ*} is the probability of the fragment starting at position *p* on *t_i_*, computed using an empirical fragment start position distribution as defined in [14]. Pr {*o|t_i_*} is the probability of obtaining a fragment aligning with the given orientation to *t_i_*. This is determined by the concordance of the fragment with the user-specified library format. It is 1 if the alignment agrees with the library format and a user-defined prior value *p_ō_* otherwise. Finally, Pr {*a | f_j_, t_i_, p, o, ℓ*} is the probability of generating alignment *a* of fragment *f_j_*, given that it is drawn from *t_i_*, with orientation *o*, and starting at position *p* and is of length *ℓ*; this term is set to 1 when using quasi-mapping, and is given by equation (7) for traditional alignments. The parameters for all auxiliary models are learned during the streaming phase of the inference algorithm from the first *N*′ observations (5,000,000 by default). These auxiliary terms can then be applied to all subsequent observations.

#### Sequence-specific bias

It has been previously observed that the sequence surrounding the 5′and 3′ ends of RNA-seq fragments has an effect on the likelihood that these fragments are selected for sequencing. If not accounted for, these biases can have a substantial effect on abundance estimates and can confound downstream analyses. To learn and correct for such biases, *Salmon* adopts a modification of the model introduced by Roberts et al. [4]. A (foreground) variable-length Markov model (VLMM) is trained on sequence windows surrounding the 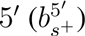 and 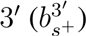 read start positions. Then, a different (background) VLMM is trained on sequence windows drawn uniformly across known transcripts, each weighted by that transcript’s abundance; the 5′ and 3′ background models are denoted as 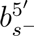 and 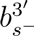 respectively.

#### Fragment GC-bias

In addition to the sequence surrounding the 5′ and 3′ ends of a fragment, it has also been observed that the GC-content of the entire fragment can play a substantial role in the likelihood that it will be selected for sequencing [5]. These biases are largely different than sequence-specific biases, and thus, accounting for both the context surrounding the fragments and the GC-content of the fragments themselves is important when one wishes to learn and correct for some of the most prevalent types of bias *in silico*. To account for fragment GC-bias, *Salmon* learns a foreground and background model of this fragment GC-bias (and defines the bias as the ratio of the score of a particular fragment under each). Our fragment GC-bias model consists of the observed distribution of sequenced fragments for every possible GC-content value (in practice, we discretize GC-content and maintain a distribution over 101 bins, for fragments with GC content ranging from 0 to 1 in increments of 0.01). The background model is trained on all possible fragments (drawn uniformly and according to the empirical fragment length distribution) across known transcripts, with each fragment weighted by that transcript’s abundance. The foreground and background fragment GC-bias models are denoted as *b*_*gc*^+^_ and *b*_*gc*^−^_ respectively. Additionally, we note that sequence-specific and fragment GC biases do seem to display a conditional dependence. To account for this, *Salmon* learns 3 different bias models, each conditioned on the average GC content of the 5′ and 3′ sequence context of the fragment. A separate model is trained and applied for fragments with average GC content between [0, 0.33), [0.33, 0.66), and [0.66, 1].

#### Incorporating the bias models

These bias models are used to re-estimate the effective length of each transcript, such that a transcript’s effective length now also takes into account the likelihood of sampling each possible fragment that transcript can produce — an approach to account for bias first introduced by Roberts et al. [4]. Before learning the bias-corrected effective lengths, the offline optimization algorithm is run for a small number of rounds (10 by default) to produce estimated abundances that are used when learning the background distributions for the various bias models. For a particular transcript *t_i_*, the effective length becomes:

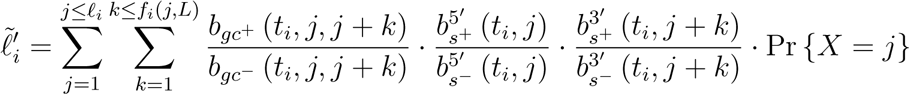

where Pr{*X* = *j*} is the probability, under the empirical fragment length distribution, of observing a fragment of length *j*, *L* is the maximum observed fragment length, *f_i_*(*j, L*) = min (*ℓ_i_* − *j* + 1, *L*), 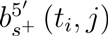 is the score given to transcript *t_i_*’s *j*^th^ position under the foreground, 5′ sequence-specific bias model 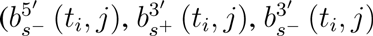 are defined similarly) and *b*_*gc*^+^_ (*t_i_, j, j* + *k*) is the score given by the foreground fragment GC-content model for the subsequence of transcript *t_i_* from position *j* to *j* + *k* (and similarly for *b*_*gc*^−^_ (*t_i_, j, j* + *k*)).

Once these bias-corrected lengths have been computed, they are used in all subsequent rounds of the offline inference phase (i.e. until the estimates of *α* — as defined in **Algorithms** — converge). Typically, the extra computational cost required to apply bias correction is rather small, and the learning and application of these bias weights is parallelized in *Salmon*. However, both the memory and time requirements of bias correction can be adjusted by the user to trade-off time and space with model fidelity. To make the computation of GC-fractions efficient for arbitrary fragments from the transcriptome, *Salmon* computes and stores the cumulative GC count for each transcript. To reduce memory consumption, this cumulative count can be sampled using the `--gcSizeSamp`. This will increase the time required to compute the GC-fraction for each fragment by a constant factor. Similarly, when attempting to determine the effective length of a transcript, *Salmon* will evaluate the contribution of all fragments longer than the shortest 0.5% and shorter than the longest 0.5% of the full empirical fragment length distribution, that could derive from this transcript. The program option `--biasSpeedSamp` will instead sample fragment lengths at a user-defined factor, speeding up the computation of bias-corrected effective lengths by this factor, but coarsening the model in the process. All results reported in this manuscript where bias correction was included were run without either of these sampling options (i.e. using the full-fidelity model).

### Alignment model

When *Salmon* is given read alignments as input, it can learn and apply a model of read alignments to help assess the probability that a fragment originated from a particular locus. Specifically, *Salmon*’s alignment model is a spatially varying first-order Markov model over the set of CIGAR symbols and nucleotides. To account for the fact that substitution and indel rates can vary spatially over the length of a read, we partition each read into a fixed number of bins (4 by default) and learn a separate model for each of these bins. This allows us to learn spatially varying effects without making the model itself too large (as if, for example, we had attempted to learn a separate model for each position in the read). Given the `CIGAR` string *s* = *s*_0_, …, *s*_|*s*|_ for an alignment *a*, we compute the probability of *a* as:

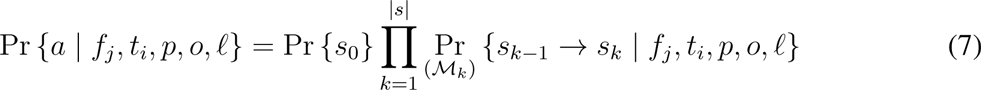

where Pr {*s*_0_} is the start probability and Pr_(*ℳ_k_*)_{·} is the transition probability under the model at the *k*^th^ position of the read (i.e., in the bin corresponding to position *k*). To compute these probabilities, *Salmon* parses the `CIGAR` string *s* and moves appropriately along both the fragment *f_j_* and the reference transcript *t_i_*, and computes the probability of transitioning to the next observed state in the alignment (a tuple consisting of the CIGAR operation, and the nucleotides in the fragment and reference) given the current state of the model. The parameters of this Markov model are learned from sampled alignments in the online phase of the algorithm (see **Algorithm 1**). When quasi-mapping is used instead of user-provided alignments, the probability of the “alignment” is not taken into account (i.e. Pr {*a | f_j_, t_i_, p, o, ℓ*} is set to 1 for each mapping).

## Algorithms

*Salmon* consists of three components: a lightweight-mapping model, an online phase that estimates initial expression levels and model parameters and constructs equivalence classes over the input fragments, and an offline phase that refines these expression estimates. The online and offline phases together optimize the estimates of *α* which is a vector of weighted estimates of read counts. Each method can compute *η* directly from these parameters.

The online phase uses a variant of stochastic, collapsed variational Bayesian inference [22]. The offline phase applies either a standard EM algorithm, or a variational Bayesian EM algorithm [21] over a reduced representation of the data represented by the equivalence classes until a data-dependent convergence criterion is satisfied. An overview of our method is given in Supplementary Fig. 1, and we describe each component in more detail below.

### Online phase

The online phase of *Salmon* attempts to solve the variational Bayesian inference problem described in **Objectives and models for abundance estimation**, and optimizes a collapsed variational objective function [19] using a variant of stochastic collapsed Variational Bayesian inference [22]. The inference procedure is a streaming algorithm, similar to [15], but it updates estimated read counts after every small group *B^τ^* (called a mini-batch) of observations, and processing of mini-batches is done asynchronously and in parallel. The pseudo-code for the algorithm is given in **Algorithm 1**.

#### Algorithm 1 Laissez-faire SCVB0

~~~
1: **while** *B^τ^* ← pop(work-queue) **do**
2:  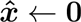
3:  **for** read *r* ∈ *B^τ^* **do**
4:    ***x* ← 0**
5:    **for** alignment *a* of *r* **do**
6:      *y* ← the transcript involved in alignment *a*
7:      *x_y_* ← *x_y_* + *α_y_* · Pr {*a | y*} ▹ Add *a*’s contribution to the local weight for transcript *y*
8:    **end for**               ▹ Normalize the contributions for all alignments of *r*
9:    **for** alignment *a* of *r* **do**
10:     *y* ← the transcript involved in alignment *a*
11:     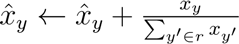
12:    **end for**
13:    Sample *a* ∈ *r* and update auxiliary models using *a*
14:  **end for**
15:  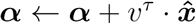 ▹ Update the global weights with local observations from *B^τ^*
16: **end while**
~~~

The observation weight for mini-batch *B^τ^*, *υ^τ^*, in line 15 of **Algorithm 1** is an increasing sequence sequence in *τ*, and is set, as in [15], to adhere to the Robbins-Monroe conditions. Here, the *α* represent the (weighted) estimated counts of fragments originating from each transcript. Using this method, the expected value of *η* can be computed directly from *α* using equation (16). We employ a *weak* Dirichlet conjugate-prior with 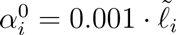 for all 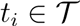. As outlined in [22], the SCVB0 inference algorithm is similar to variants of the online-EM [23] algorithm with a modified prior. The procedure in **Algorithm 1** is run independently by as many worker threads as the user has specified. The threads share a single work-queue upon which a parsing thread places mini-batches of alignment groups. An alignment group is simply the collection of all alignments (i.e. all multi-mapping locations) for a particular read. The mini-batch itself consists of a collection of some small, fixed number of alignment groups (1,000 by default). Each worker thread processes one alignment group at a time, using the current weights of each transcript and the current auxiliary parameters to estimate the probability that a read came from each potential transcript of origin. The processing of mini-batches occurs in parallel, so that very little synchronization is required, only an atomic compare-and-swap loop to update the global transcript weights at the end of processing of each mini-batch — hence the moniker laissez-faire in the label of **Algorithm 1**. This lack of synchronization means that when estimating *x_y_*, we cannot be certain that the most up-to-date values of *α* are being used. However, due to the stochastic and additive nature of the updates, this has little-to-no detrimental effect [24]. The inference procedure itself is generic over the type of alignments being processed; they may be either regular alignments (e.g. coming from a `bam` file), or quasi-mappings computed from the raw reads (e.g. coming from `FASTA/Q` files). After the entire mini-batch has been processed, the global weights for each transcript are updated. These updates are *sparse*; i.e. only transcripts that appeared in some alignment in mini-batch *B^τ^* will have their global weight updated after *B^τ^* has been processed. This ensures, as in [15], that updates to the parameters *α* can be performed efficiently.

### Equivalence classes

During its online phase, in addition to performing streaming inference of transcript abundances, *Salmon* also constructs a highly-reduced representation of the sequencing experiment. Specifically, *Salmon* constructs “rich” equivalence classes over all of the sequenced fragments. Collapsing fragments into equivalence classes is a well-established idea in the transcript quantification literature, and numerous different notions of equivalence classes have been previously introduced, and shown to greatly reduce the time required to perform iterative optimization such as that described in **Offline phase**. For example, Salzman et al. [25] first introduced the notion of factorizing the likelihood function to speed up inference by collapsing fragments that align to the same exons or exon junctions (as determined by a provided annotation) into equivalence classes. Simlarly, Nicolae et al.[26] used equivalence classes over fragments to reduce memory usage and speed up inference — they define as equivalent any pair of fragments that align to the same set of transcripts and whose compatibility weights (i.e. conditional probabilities) with respect to those transcripts are proportional. Patro et al. [9] define equivalence classes over k-mers, treating as equivalent any k-mers that appear in the same set of transcripts at the same frequency, and use this factorization of the likelihood function to speed up optimization. Bray et al. [10] define equivalence classes over fragments, and define as equivalent any fragments that pseuoalign to the same set of transcripts — this is similar to the notion adopted by Nicolae et al., except that no restriction is placed on the proportionality of compatibility weights (since these are not computed).

To compute equivalence classes, we define an equivalence relation ∼ over fragments. Let 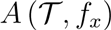 denote the set of quasi-mappings (or alignments) of *f_x_* to the transcriptome 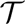, and let 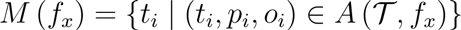 be the set of transcripts to which *f_x_* maps according to 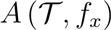. We say *f_x_* ∼ *f_y_* if and only if *M*(*f_x_*) = *M*(*f_y_*). Fragments which are equivalent are grouped together for the purpose of inference. *Salmon* builds up a set of fragment-level equivalence classes by maintaining an efficient concurrent cuckoo hash map [27]. To construct this map, we associate each fragment *f_x_* with *t^x^* = *M*(*f_x_*), which we will call the label of the fragment. Then, we query the hash map for *t^x^*. If this key is not in the map, we create a new equivalence class with this label, and set its count to 1. Otherwise, we increment the count of the equivalence class that we find in the map with this label. The efficient, concurrent nature of the data structure means that many threads can simultaneously query and write to the map while encountering very little contention. Each key in the hash map is associated with a value that we call a “rich” equivalence class. For each equivalence class 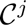, we retain a count 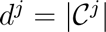, which is the total number of fragments contained within this class. We also maintain, for each class, a weight vector *w^j^*. The entries of this vector are in one-to-one correspondence with transcripts *i* in the label of this equivalence class such that

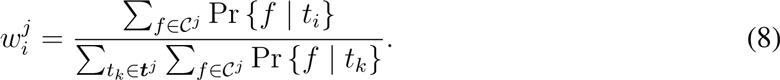

That is, 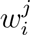 is the average conditional probability of observing a fragment from 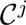 given *t_i_* over all fragments in this equivalence class. Though the likelihood function over equivalence classes that considers these weights (Equation (10)) is no longer exactly equivalent to the likelihood defined over all fragments (Equation (9)), these weights nonetheless allow us to take into consideration the conditional probabilities specified in the full model, without having to continuously reconsider each of the fragments in *ℱ*. There is a spectrum of possible representations of “rich” equivalence classes. This spectrum spans from notion adopted here, which collapses all conditional probabilities into a single aggregate scalar, to an approach that clusters together fragments based not only on the transcripts to which they match, but on the vector of normalized conditional probabilities for each of these transcripts. The former approach represents a more coarse-grained approximate factorization of the likelihoodod function while the latter represents a more fine-grained approximation. We believe that studying how these different notions of equivalence classes affect the factorization of the likelihood function, and hence its optimization, is an interesting direction for future work.

### Offline phase

In its offline phase, which follows the online phase, *Salmon* uses the “rich” equivalence classes learned during the online phase to refine the inference. Given the set 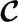 of rich equivalence classes of fragments, we can use an expectation maximization (EM) algorithm to optimize the likelihood of the parameters given the data. The abundances *η* can be computed directly from *α*, and we compute maximum likelihood estimates of these parameters which represent the estimated counts (i.e. number of fragments) deriving from each transcript, where:

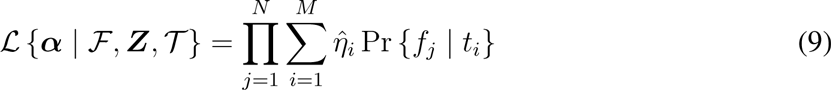

and 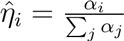. If we write this same likelihood in terms of the equivalence classes 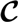, we have:

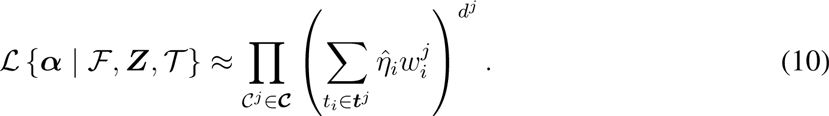

#### EM update rule

This likelihood, and hence that represented in equation (9), can then be optimized by applying the following update equation iteratively

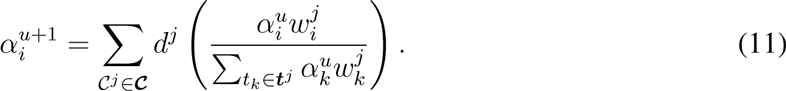

We apply this update equation until the maximum relative difference in the *α* parameters satisfies:

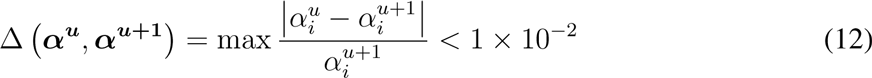

for all 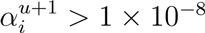. Let *α^′^* be the estimates after having achieved convergence. We can then estimate *η_i_* by 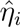, where:

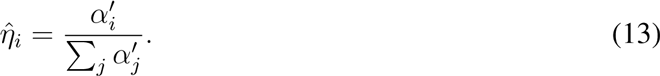

#### Variational Bayes optimization

Instead of the standard EM updates of equation (11), we can, optionally, perform Variational Bayesian optimization by applying VBEM updates as in [20], but adapted to be with respect to the equivalence classes:

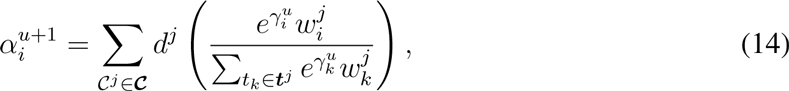

where:

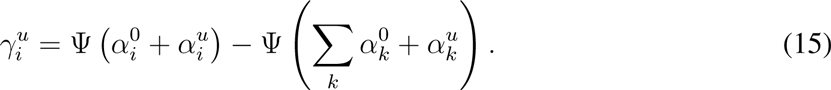

Here, Ψ(*·*) is the digamma function, and, upon convergence of the parameters, we can obtain an estimate of the expected value of the posterior nucleotide fractions as:

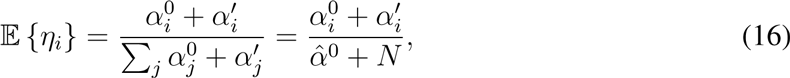

where 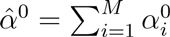. Variational Bayesian optimization in the offline-phase of *Salmon* is selected by passing the `--useVBOpt` flag to the *Salmon* quant command.

### Sampling from the posterior

After the convergence of the parameter estimates has been achieved in the offline phase, it is possible to draw samples from the posterior distribution using collapsed, blockwise Gibbs sampling over the equivalence classes. Samples can be drawn by iterating over the equivalence classes, and re-sampling assignments for some fraction of fragments in each class according to the multinomial distribution defined by holding the assignments for all other fragments fixed. Many samples can be drawn quickly, since many Gibbs chains can be run in parallel. Further, due to the accuracy of the preceding inference, the chains begin sampling from a relatively high probability position in the latent variable space almost immediately. These posterior samples can be used to obtain estimates for quantities of interest about the posterior distribution, such as its variance, or to produce credible intervals. When *Salmon* is passed the `--numGibbsSamples` option, it will draw a number of posterior samples that is provided to this option.

Additionally, inspired by *kallisto* [10], *Salmon* also provides the ability to draw bootstrap samples, which is an alternative way to assign confidence to the estimates returned by the main inference algorithm. Bootstrap samples can be drawn by passing the --numBootstraps option to *Salmon* with the argument determining the number of bootstraps to perform. The bootstrap sampling process works by sampling (with replacement) counts for each equivlalence class, and then re-running the offline inference procedure (either the EM or VBEM algorithm) for each bootstrap sample.

## Validation

### Metrics for accuracy

Throughout this paper, we use several different different metrics to summarize the agreement of the estimated TPM for each transcript with the TPM computed from simulated counts. While most of these metrics are commonly used and self-explanatory, we here describe the computation of the mean absolute relative difference (MARD), which is, less common than some of the other metrics.

The MARD is computed using the absolute relative difference ARD_*i*_ for each transcript *i*:

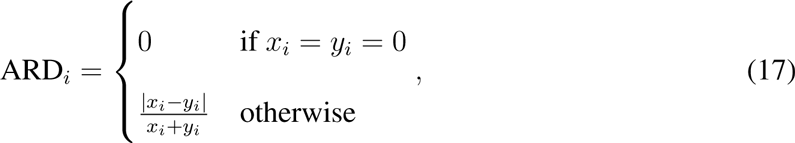

where *x_i_* is the true value of the TPM, and *y_i_* is the estimated TPM. The relative difference is bounded above by 1, and takes on a value of 0 whenever the prediction perfectly matches the truth. To compute the mean absolute relative difference, we simply take 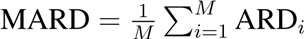. We note that *Salmon* and *kallisto*, by default, truncate very tiny expression values to 0. For example, any transcript estimated to produce < 1 × 10^−8^ reads is assigned an estimated read count of 0 (which, likewise, affects the TPM estimates). However, *eXpress* does not perform such a truncation, and very small, non-zero values may have a negative effect on the MARD metric. To mitigate such effects, we first truncate to 0 all TPMs less than 0.01 before computing the MARDs.

### Ground truth simulated data

To assess accuracy in a situation where the true expression levels are known, we generate synthetic data sets using both *Polyester* [13] and *RSEM-sim* [14].

#### *RSEM-sim* simulations

To generate data with *RSEM-sim*, we follow the procedure used in [10]. *RSEM* was run on sample `NA12716_7` from the GEUVADIS RNA-seq data to learn model parameters and estimate true expression, and the learned model was then used to generate 20 different simulated datasets, each consisting of 30 million 75 bp paired-end reads.

#### *Polyester* simulations

In addition to the ability to generate reads, *Polyester* allows simulating experiments with differential transcript expression and biological variability. Thus, we can assess not only the accuracy of the resulting estimates, but also how these estimates would perform in a typical downstream analysis task like differential expression testing.

The *Polyester* simulation of an RNA-seq experiment with empirically-derived fragment GC bias was created as follows: The transcript abundance quantifications from *RSEM* run on `NA12716_7` of the GEUVADIS RNA-seq data [11] were summed to the gene-level using version 75 of the Ensembl gene annotation for GRCh37. Subsequently, whole-transcriptome simulation was carried out using *Polyester*. Abundance (TPMs) was allocated to isoforms within a gene randomly using the following rule: for genes with two isoforms, TPMs were either (i) split according to a flat Dirichlet distribution (*α* = (1, 1)) or (ii) attributed to a single isoform. The choice of (i) vs (ii) was decided by a Bernoulli trial with probability 0.5. For genes with three or more isoforms, TPMs were either (i) split among three randomly chosen isoforms according to a flat Dirichlet distribution (*α* = (1, 1, 1)) or (ii) attributed to a single isoform. Again, (i) vs (ii) was decided by a Bernoulli trial with probability 0.5. The choice of distributing expression among three isoforms was motivated by exploratory data analysis of estimated transcript abundance revealing that for most genes nearly all of expression was concentrated in the first three isoforms for genes with four or more isoforms.

Expected counts for each transcript were then generated according to the transcript-level TPMs, multiplied by the transcript lengths. 40 million 100bp paired-end reads were simulated using the *Polyester* software for each of 16 samples, and 10% of transcripts were chosen to be differentially expressed across an 8 vs 8 sample split. The fold change was chosen to be either 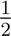 or 2 with probability of 0.5. Fragments were down-sampled with Bernoulli trials according to an empirically-derived fragment GC content dependence estimated with *alpine* [5] on RNA-seq samples from the GEUVADIS project. The first 8 GEUVADIS samples exhibited weak GC content dependence while the last 8 samples exhibited more severe fragment-level GC bias. Paired-end fragments were then shuffled before being supplied to transcript abundance quantifiers. Estimated expression was compared to true expression calculated on transcript counts (before these counts were down-sampled according to the empirically-derived fragment GC bias curve), divided by effective transcript length and scaled to TPM. Global differences across condition for all methods were removed using a scaling factor per condition. Differences across condition for the different methods’ quantifications were tested using a t-test of log_2_(**TPM** + 1).

## Software versions and options

All tests were performed with *eXpress* v1.5.1, *kallisto* v0.43.0, *Salmon* v0.7.1 and Bowtie2 v2.2.4. Reads were aligned with Bowtie2 using the parameters `--no-discordant -k 200`, and `-p` to set the number of threads. On the *RSEM-sim* data, all methods were run *without* bias correction. On all other datasests, methods were run with bias correction unless otherwise noted. Additionally, on the *Polyester* simulated data, *Salmon* was run with the option `--noBiasLengthThreshold`, which allows bias correction, even for very short transcripts, since we were most interested in assessing the maximum sensitivity of the model.

### GEUVADIS data

The analyses presented in Fig. 1d, Supplementary Table 1 and Supplementary Fig. 5 were carried out on a subset of 30 samples from the publicly-available GEUVADIS [11] data. The accesssions used and the information about the center at which the libraries were prepared and sequenced is recorded in Supplementary Table 2. All methods were run with bias correction enabled, using a transcriptome built with the RefSeq gene annotation file and the genome FASTA contained within the hg19 Illumina iGenome, in order to allow for comparison with the results in [5]. For each transcript, a t-test was performed, comparing log_2_(**TPM**+1) from 15 samples from one sequencing center against 15 samples from another sequencing center. Because the samples are from the same human population, it is expected that there would be few to no true differences in transcript abundance produced by this comparison. *P* values were then adjusted using the method of Benjamini-Hochberg, over the transcripts with mean **TPM** > 0.1. The number of positives for given false discovery rates was then reported for each method, by taking the number of transcripts with adjusted *p* value less than a given threshold.

### SEQC data

The consistency analysis presented in Supplementary Fig. 2 was carried out on a subset of the publicly-available SEQC [28] data. Specifically, the accessions used, along with the corresponding information about the center at which they were sequences is recorded in Supplementary Table 3. For each sample, “same center” comparisons were made between all unique pairs of replicates labeled as coming from the same sequencing center, while “different center” comparisons were made between all unique pairs of replicates labeled as coming from different centers (“Center” column of Supplementary Table 3).

## Supplementary Material

**Supplementary Figure 1:**
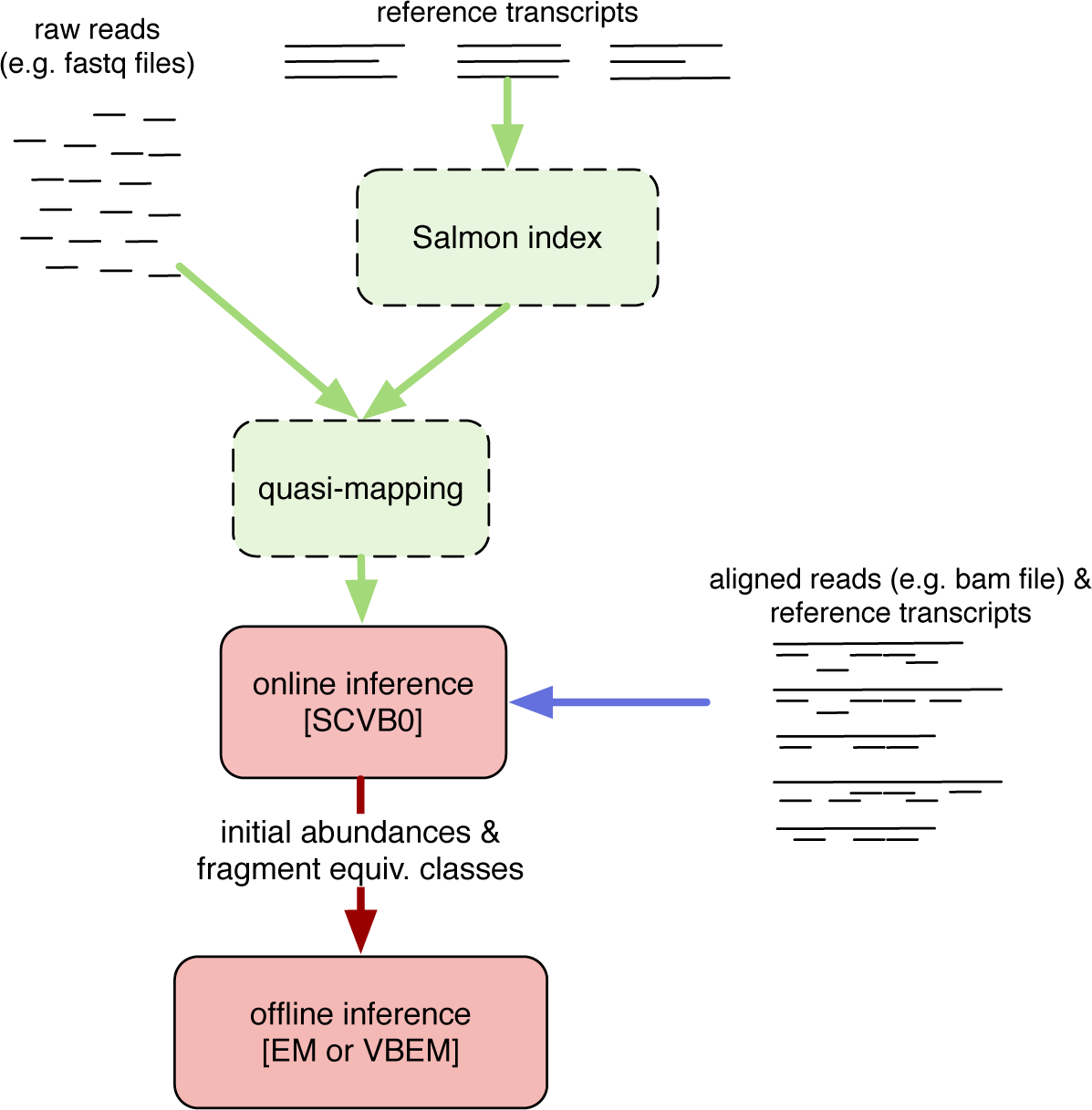
Overview of Salmon’s method and components. Supplementary Figure 1: Overview of *Salmon*’s method and components. *Salmon* accepts either raw (green arrows) or aligned reads (blue arrow) as input, performs an online inference when processing fragments or alignments, builds equivalence classes over these fragments and subsequently refines abundance estimates using an offline inference algorithm on a reduced representation of the data.

**Supplementary Figure 2:**
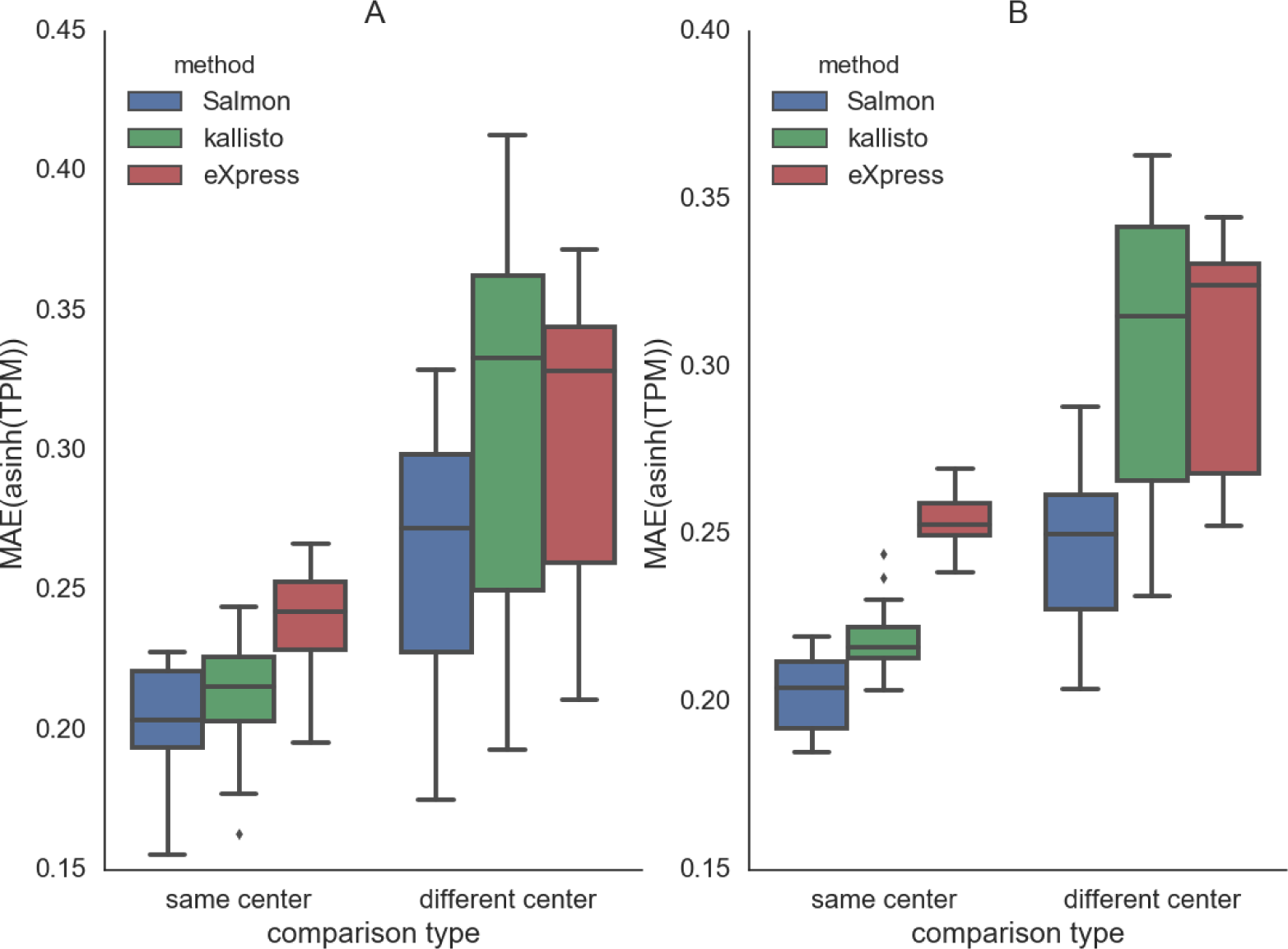
Consistency of estimates on SEQC data within and between centers. Supplementary Figure 2: The distribution of the mean absolute error of (inverse hyperbolic sine-transformed) TPMs between different replicates of data from the SEQC [28] study. The A sample corresponds to universal human reference tissue (UHRR) and the B sample corresponds to human brain tissue (HBRR). When comparing the replicates that were sequenced at different centers, the inter-replicate distances are larger. However, we observe that *Salmon*’s bias correction methodology results in improved consistency (i.e. reduced distance) compared to the estimates produced by other methods, especially when comparing replicates sequenced at different centers, where we expect the effects of bias to be more pronounced.

**Supplementary Figure 3:**
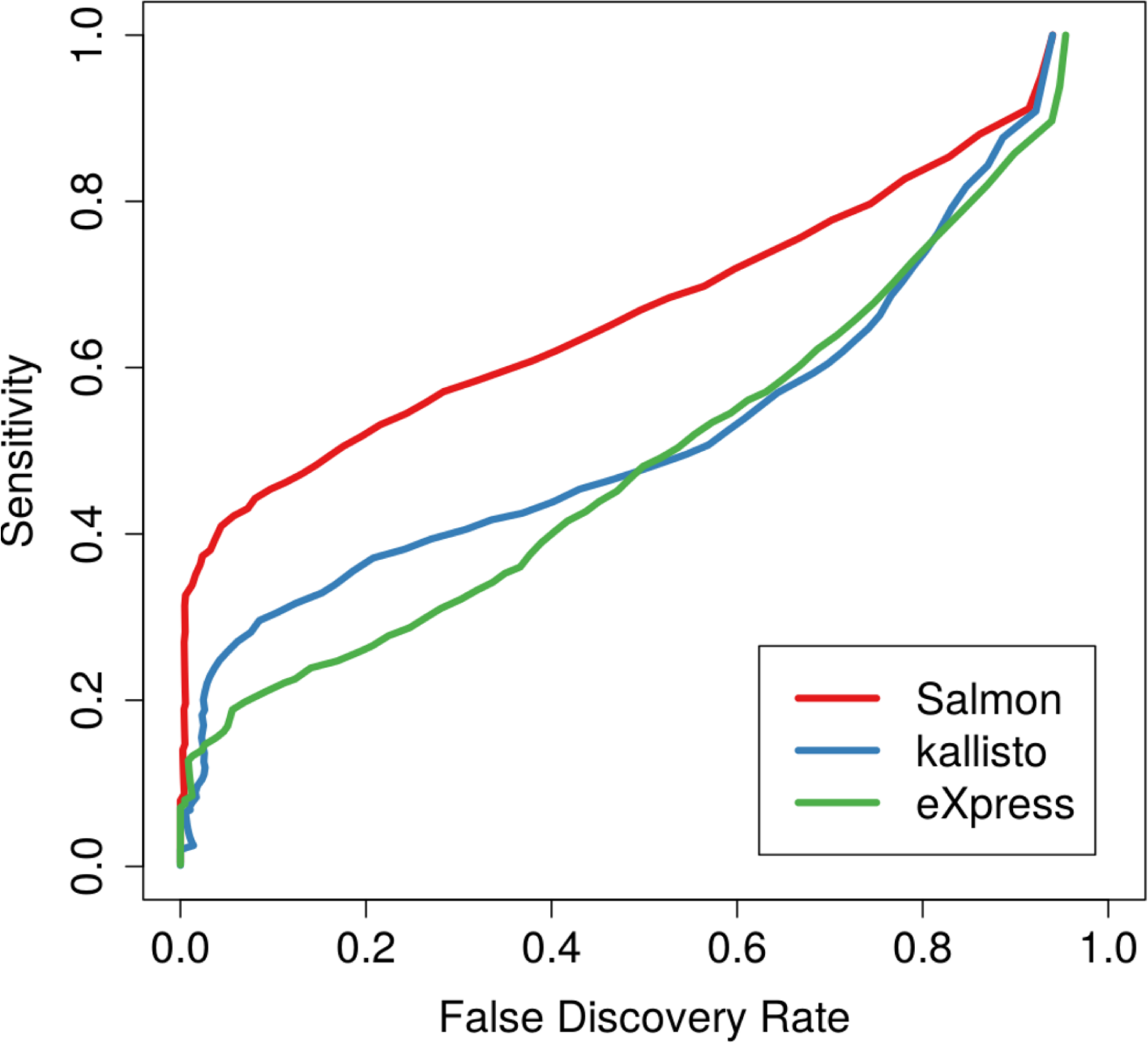
FDR vs. sensitivity on *Polyester* simulated data. Supplementary Figure 3: The false discovery rate (FDR) vs. the sensitivity of *Salmon*, *kallisto* and *eXpress* on *Polyester* simulated RNA-seq data using empirically-derived fragment GC bias profiles. All methods were run with bias-correction enabled, but only *Salmon*’s model incorporates corrections for fragment GC bias. This leads to a large improvement in sensitivity at almost every FDR value.

**Supplementary Figure 4:**
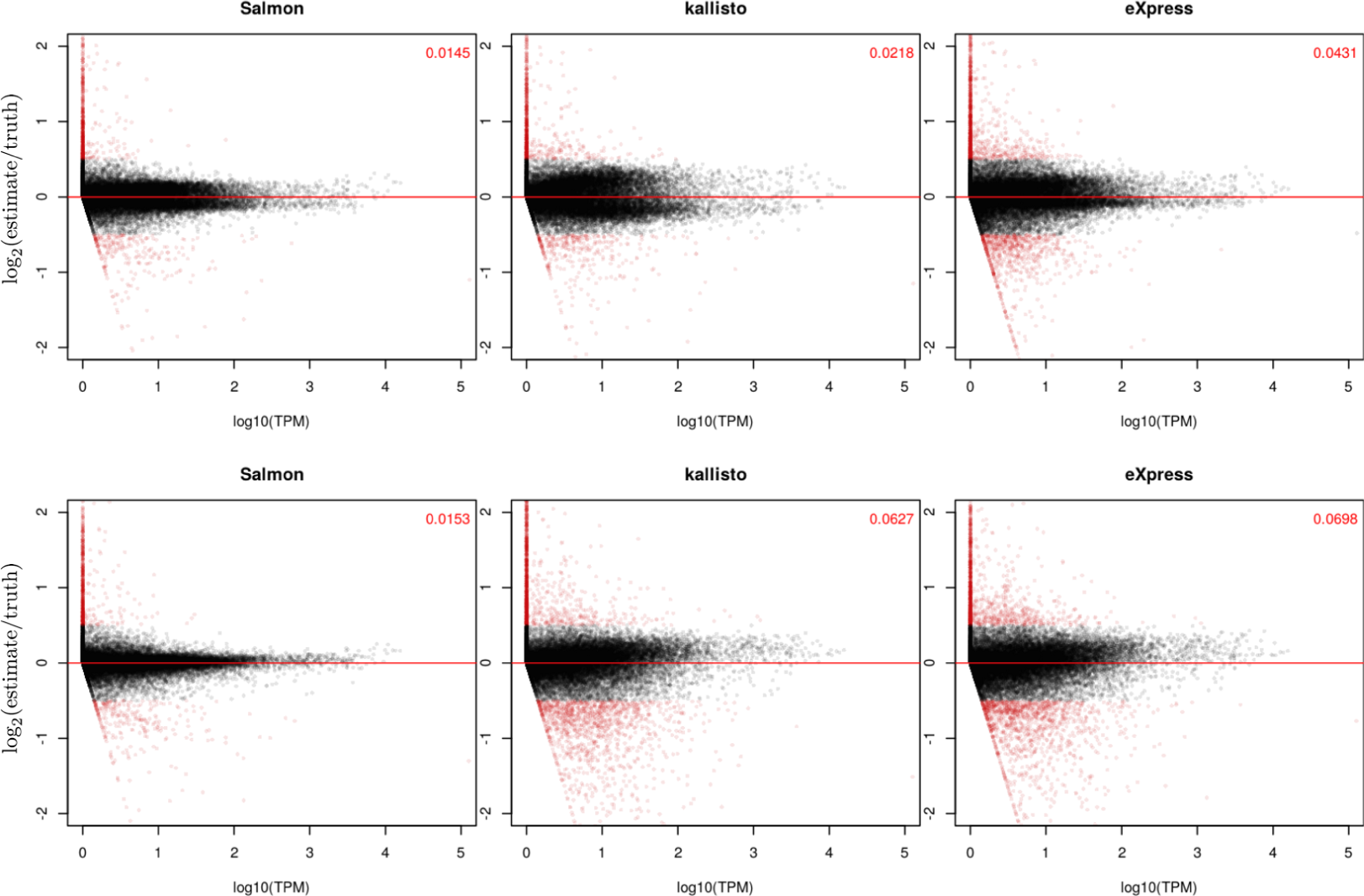
Abundance vs. fold change accuracy on *Polyester* simulated data. Supplementary Figure 4: The log_2_ fold change between the estimated and true abundances as a function of the true abundance (measured in TPM), for all 3 methods and for all replicates of both simulated “conditions” (each row displays points from all samples within a given condition). The top row corresponds to the 8 samples simulated from the data showing the weak fragment GC content bias, while the bottom row corresponds to the 8 samples simulated from the data showing the stronger fragment GC content bias. Points with an estimated log_2_ fold change of > 0.5 or < −0.5 are colored red. The fraction of red points appears in the upper right-hand corner of each plot. *Salmon* consistently demonstrates log fold changes closer to 0 than either *kallisto* or *eXpress*, across most of the range of expression.

**Supplementary Table 1:** Gene Level GEUVADIS DE. Supplementary Table 1: The number of genes identified as differentially expressed at a target FDR of 1% for two groups of GEUVADIS samples. The contrast between samples is a technical confound, and we expect little-to-no true DE. Gene-level TPM was computed by summing the TPM of the isoforms, and differential testing was performed as described in Methods.

| *Salmon* | *kallisto* | *eXpress* |
| --- | --- | --- |
| 455 | 1200 | 1582 |

**Supplementary Figure 5:**
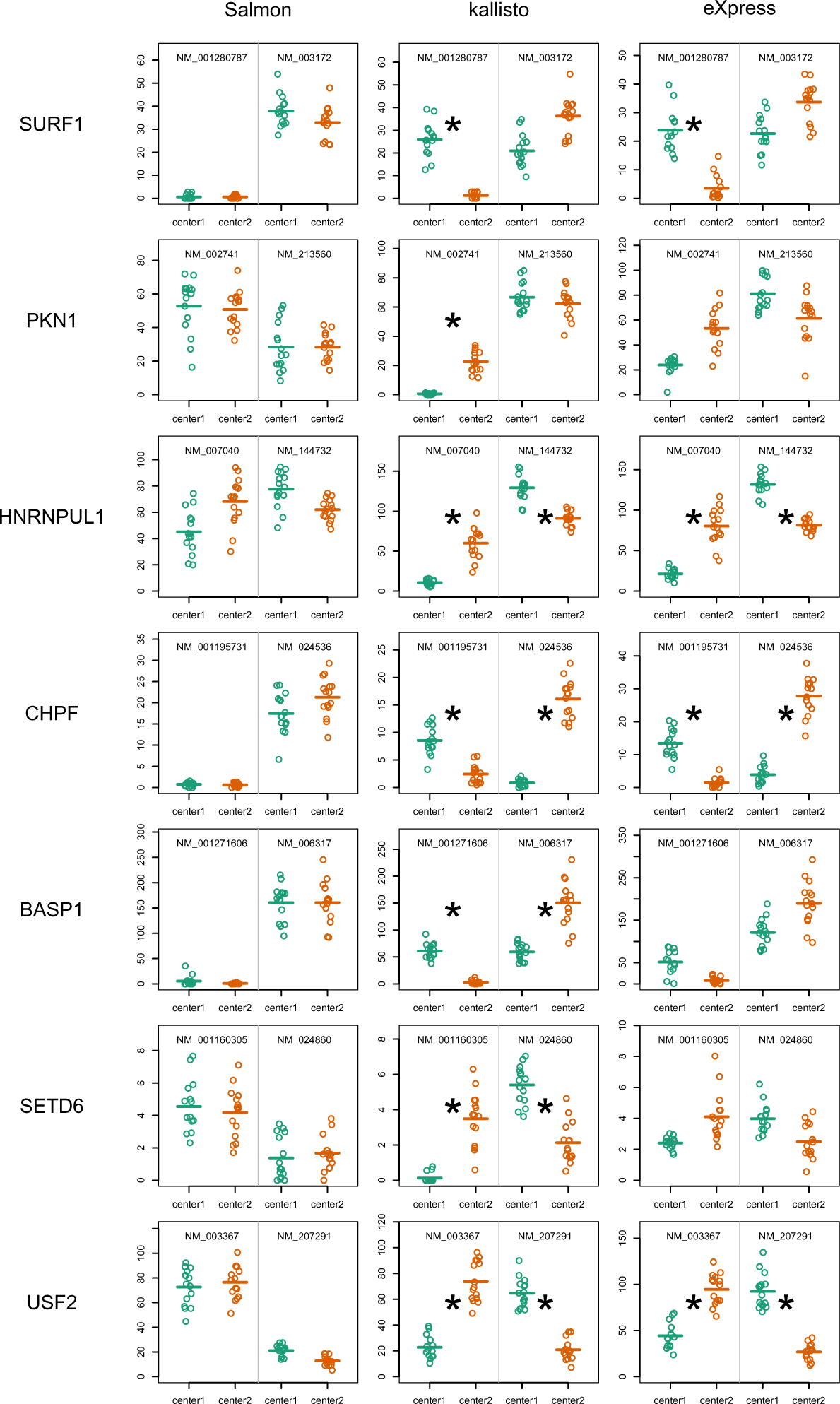
*Salmon* reduces false isoform switching. Supplementary Figure 5: Transcripts demonstrating dominant isoform switching that results from technical bias. In the quantification estimates computed using *kallisto* and *eXpress*, these two-isoform genes show a change in the dominant isoform between conditions (an asterisk denotes a t-test on log_2_(TPM+1) with *p* < 1 × 10^−6^). However, *Salmon* directly corrects for technical biases that appear to underlay differences across sequencing center, revealing that the dominant isoform has not, in fact, switched across center.

**Supplementary Figure 6:**
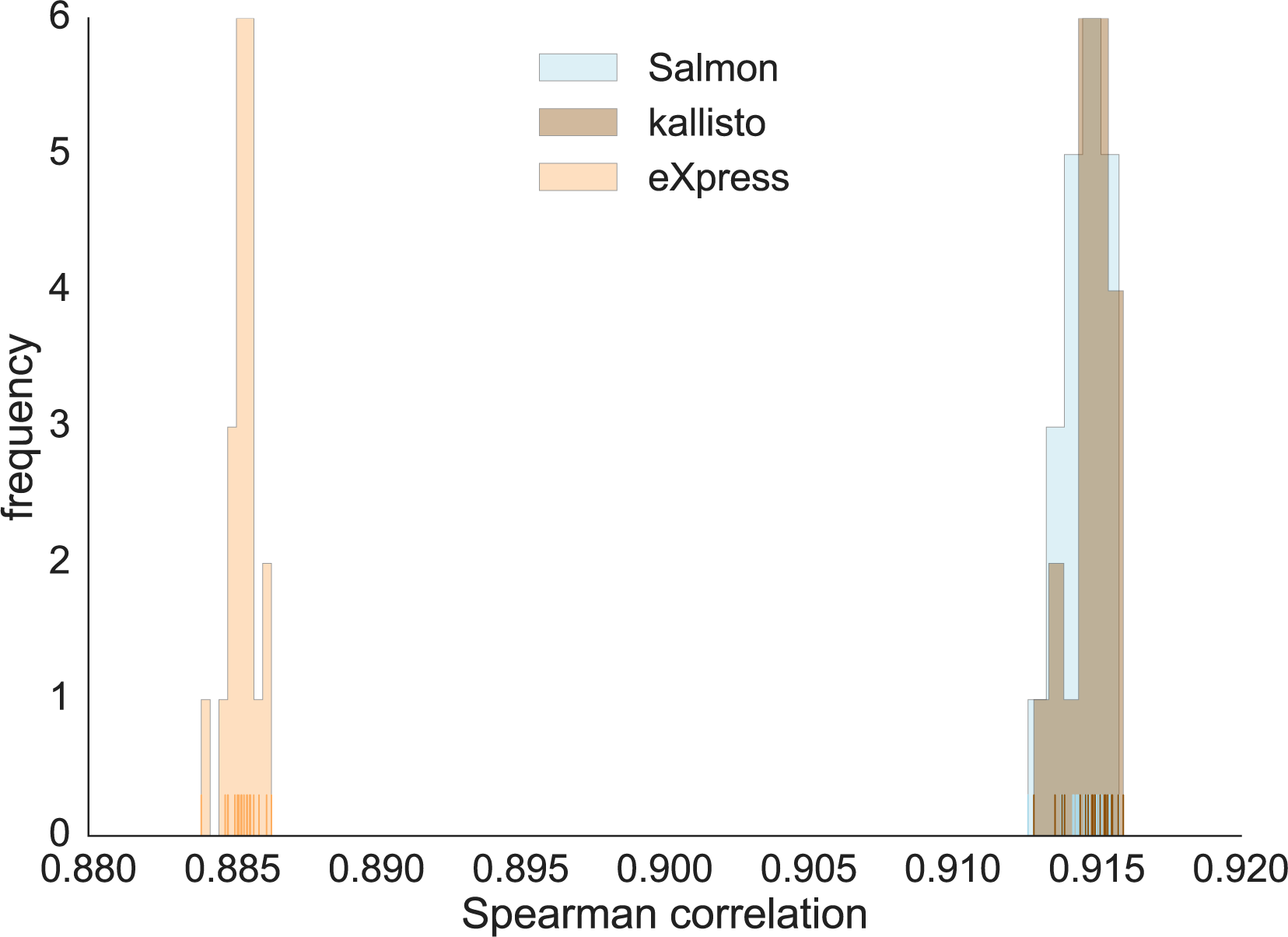
Quantification accuracy for *Salmon*, *kallisto* and *eXpress* using *RSEM-sim* data. Supplementary Figure 6: This plot shows the distribution of Spearman correlations over all 20 replicates of the *RSEM-sim* data for *Salmon*, *kallisto* and *eXpress*. *Salmon* and *kallisto* yield very similar distributions of correlations (no statistically significant difference), while both methods yeild correlations greater than that of *eXpress* (Mann-Whitney *U* test, *p* = 3.39780 × 10^−8^).

**Supplementary Table 1:**
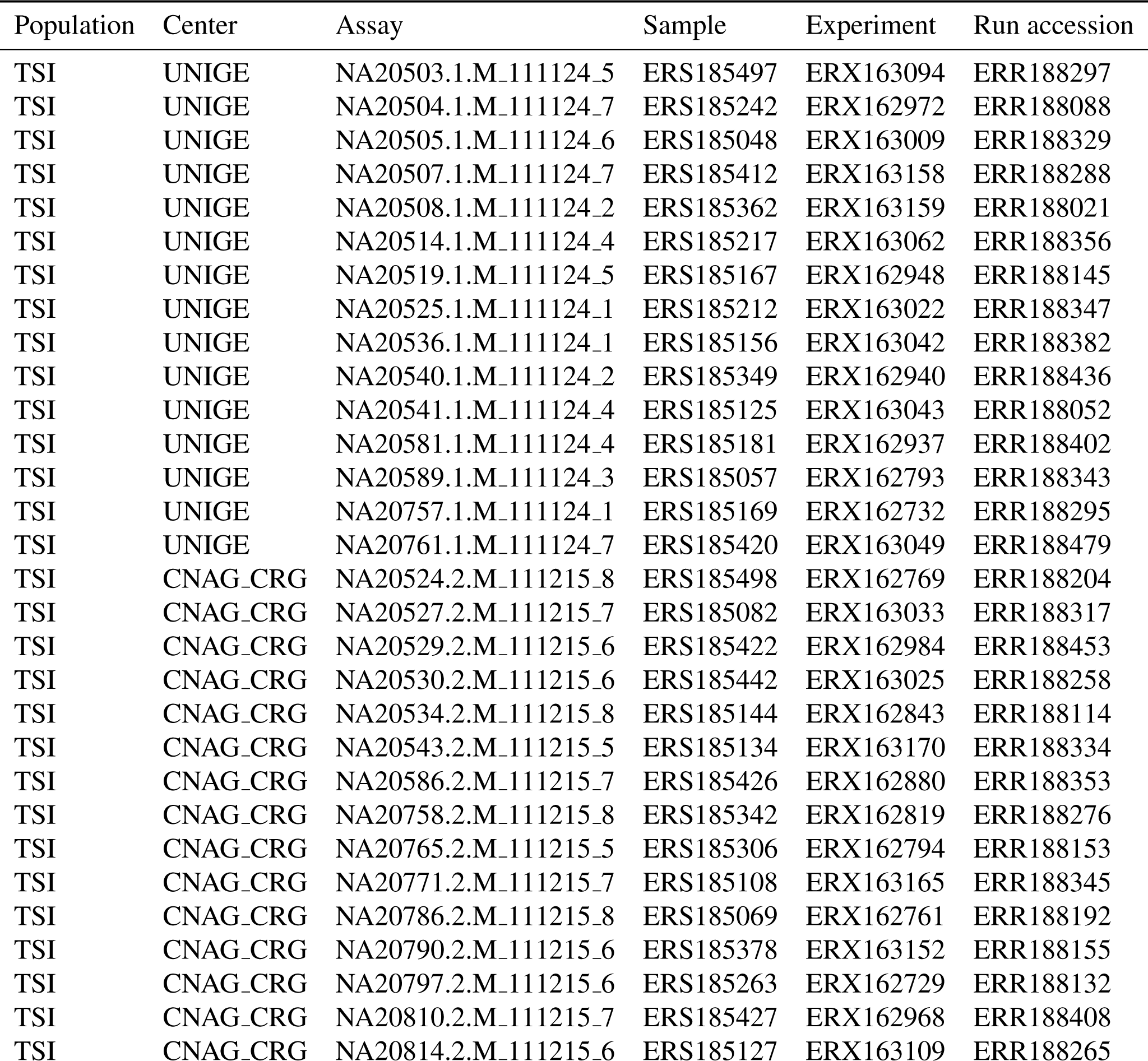
GEUVADIS samples used for specificity assessment. Supplementary Table 2: Accession information for GEUVADIS samples.

**Supplementary Table 3:**
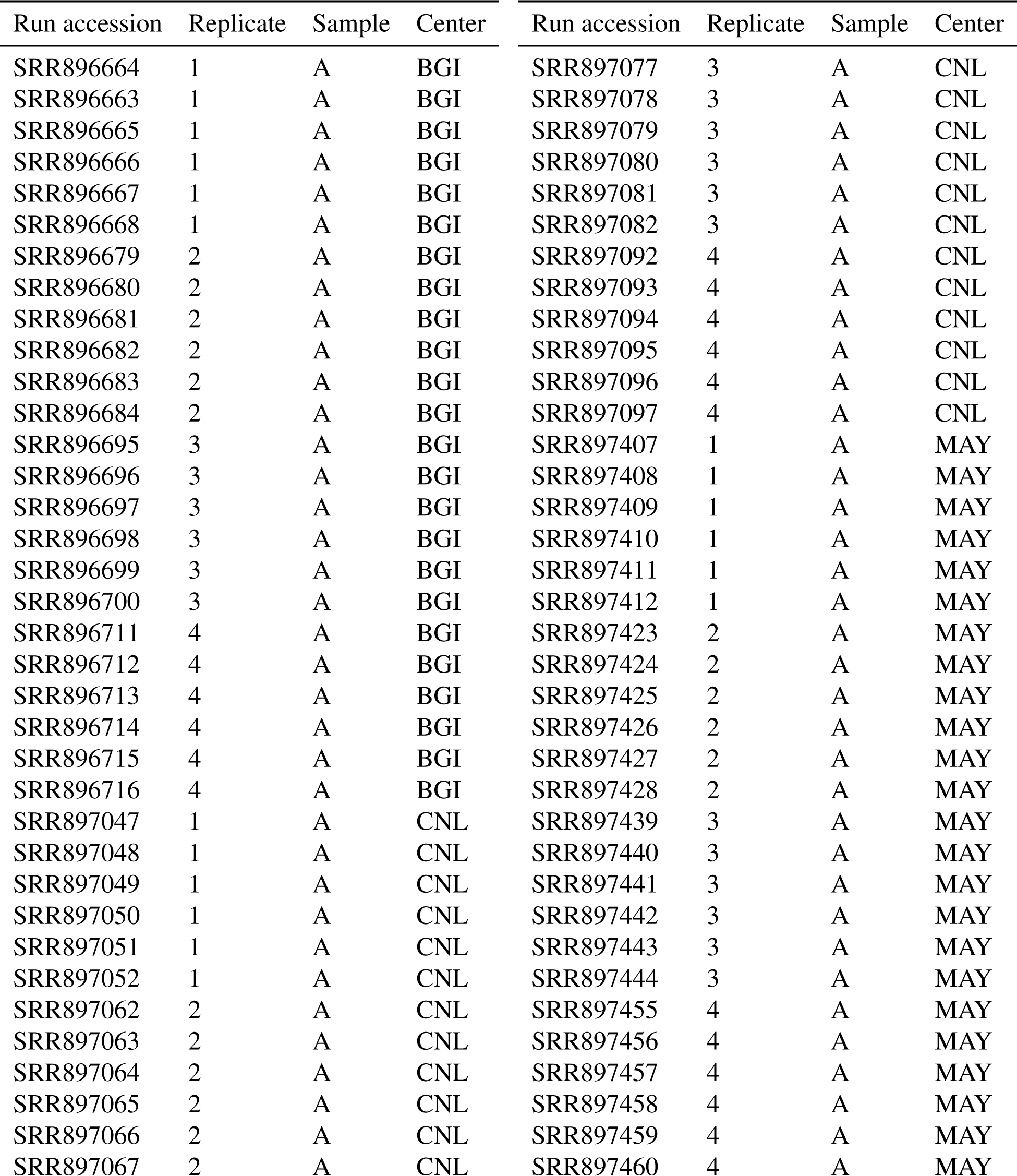
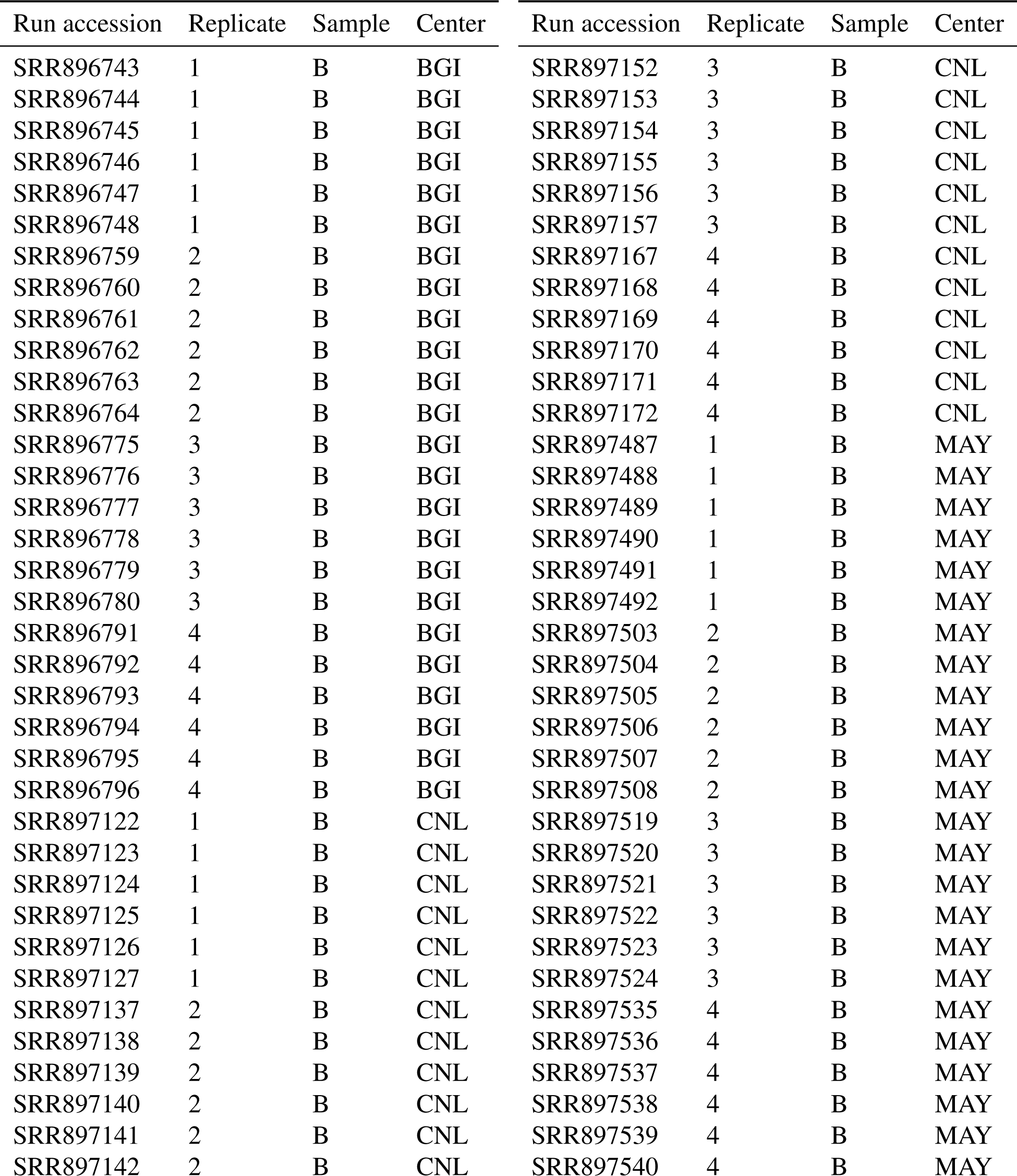
SEQC samples used for consistency assessment. Supplementary Table 3: Accession information for SEQC samples.

| Run accession | Replicate | Sample | Center | Run accession | Replicate | Sample | Center |
| --- | --- | --- | --- | --- | --- | --- | --- |
| SRR896664 | 1 | A | BGI | SRR897077 | 3 | A | CNL |
| SRR896663 | 1 | A | BGI | SRR897078 | 3 | A | CNL |
| SRR896665 | 1 | A | BGI | SRR897079 | 3 | A | CNL |
| SRR896666 | 1 | A | BGI | SRR897080 | 3 | A | CNL |
| SRR896667 | 1 | A | BGI | SRR897081 | 3 | A | CNL |
| SRR896668 | 1 | A | BGI | SRR897082 | 3 | A | CNL |
| SRR896679 | 2 | A | BGI | SRR897092 | 4 | A | CNL |
| SRR896680 | 2 | A | BGI | SRR897093 | 4 | A | CNL |
| SRR896681 | 2 | A | BGI | SRR897094 | 4 | A | CNL |
| SRR896682 | 2 | A | BGI | SRR897095 | 4 | A | CNL |
| SRR896683 | 2 | A | BGI | SRR897096 | 4 | A | CNL |
| SRR896684 | 2 | A | BGI | SRR897097 | 4 | A | CNL |
| SRR896695 | 3 | A | BGI | SRR897407 | 1 | A | MAY |
| SRR896696 | 3 | A | BGI | SRR897408 | 1 | A | MAY |
| SRR896697 | 3 | A | BGI | SRR897409 | 1 | A | MAY |
| SRR896698 | 3 | A | BGI | SRR897410 | 1 | A | MAY |
| SRR896699 | 3 | A | BGI | SRR897411 | 1 | A | MAY |
| SRR896700 | 3 | A | BGI | SRR897412 | 1 | A | MAY |
| SRR896711 | 4 | A | BGI | SRR897423 | 2 | A | MAY |
| SRR896712 | 4 | A | BGI | SRR897424 | 2 | A | MAY |
| SRR896713 | 4 | A | BGI | SRR897425 | 2 | A | MAY |
| SRR896714 | 4 | A | BGI | SRR897426 | 2 | A | MAY |
| SRR896715 | 4 | A | BGI | SRR897427 | 2 | A | MAY |
| SRR896716 | 4 | A | BGI | SRR897428 | 2 | A | MAY |
| SRR897047 | 1 | A | CNL | SRR897439 | 3 | A | MAY |
| SRR897048 | 1 | A | CNL | SRR897440 | 3 | A | MAY |
| SRR897049 | 1 | A | CNL | SRR897441 | 3 | A | MAY |
| SRR897050 | 1 | A | CNL | SRR897442 | 3 | A | MAY |
| SRR897051 | 1 | A | CNL | SRR897443 | 3 | A | MAY |
| SRR897052 | 1 | A | CNL | SRR897444 | 3 | A | MAY |
| SRR897062 | 2 | A | CNL | SRR897455 | 4 | A | MAY |
| SRR897063 | 2 | A | CNL | SRR897456 | 4 | A | MAY |
| SRR897064 | 2 | A | CNL | SRR897457 | 4 | A | MAY |
| SRR897065 | 2 | A | CNL | SRR897458 | 4 | A | MAY |
| SRR897066 | 2 | A | CNL | SRR897459 | 4 | A | MAY |
| SRR897067 | 2 | A | CNL | SRR897460 | 4 | A | MAY |
| SRR896743 | 1 | B | BGI | SRR897152 | 3 | B | CNL |
| SRR896744 | 1 | B | BGI | SRR897153 | 3 | B | CNL |
| SRR896745 | 1 | B | BGI | SRR897154 | 3 | B | CNL |
| SRR896746 | 1 | B | BGI | SRR897155 | 3 | B | CNL |
| SRR896747 | 1 | B | BGI | SRR897156 | 3 | B | CNL |
| SRR896748 | 1 | B | BGI | SRR897157 | 3 | B | CNL |
| SRR896759 | 2 | B | BGI | SRR897167 | 4 | B | CNL |
| SRR896760 | 2 | B | BGI | SRR897168 | 4 | B | CNL |
| SRR896761 | 2 | B | BGI | SRR897169 | 4 | B | CNL |
| SRR896762 | 2 | B | BGI | SRR897170 | 4 | B | CNL |
| SRR896763 | 2 | B | BGI | SRR897171 | 4 | B | CNL |
| SRR896764 | 2 | B | BGI | SRR897172 | 4 | B | CNL |
| SRR896775 | 3 | B | BGI | SRR897487 | 1 | B | MAY |
| SRR896776 | 3 | B | BGI | SRR897488 | 1 | B | MAY |
| SRR896777 | 3 | B | BGI | SRR897489 | 1 | B | MAY |
| SRR896778 | 3 | B | BGI | SRR897490 | 1 | B | MAY |
| SRR896779 | 3 | B | BGI | SRR897491 | 1 | B | MAY |
| SRR896780 | 3 | B | BGI | SRR897492 | 1 | B | MAY |
| SRR896791 | 4 | B | BGI | SRR897503 | 2 | B | MAY |
| SRR896792 | 4 | B | BGI | SRR897504 | 2 | B | MAY |
| SRR896793 | 4 | B | BGI | SRR897505 | 2 | B | MAY |
| SRR896794 | 4 | B | BGI | SRR897506 | 2 | B | MAY |
| SRR896795 | 4 | B | BGI | SRR897507 | 2 | B | MAY |
| SRR896796 | 4 | B | BGI | SRR897508 | 2 | B | MAY |
| SRR897122 | 1 | B | CNL | SRR897519 | 3 | B | MAY |
| SRR897123 | 1 | B | CNL | SRR897520 | 3 | B | MAY |
| SRR897124 | 1 | B | CNL | SRR897521 | 3 | B | MAY |
| SRR897125 | 1 | B | CNL | SRR897522 | 3 | B | MAY |
| SRR897126 | 1 | B | CNL | SRR897523 | 3 | B | MAY |
| SRR897127 | 1 | B | CNL | SRR897524 | 3 | B | MAY |
| SRR897137 | 2 | B | CNL | SRR897535 | 4 | B | MAY |
| SRR897138 | 2 | B | CNL | SRR897536 | 4 | B | MAY |
| SRR897139 | 2 | B | CNL | SRR897537 | 4 | B | MAY |
| SRR897140 | 2 | B | CNL | SRR897538 | 4 | B | MAY |
| SRR897141 | 2 | B | CNL | SRR897539 | 4 | B | MAY |
| SRR897142 | 2 | B | CNL | SRR897540 | 4 | B | MAY |

